# *De novo*-designed minibinders expand the synthetic biology sensing repertoire

**DOI:** 10.1101/2024.01.12.575267

**Authors:** Zara Y. Weinberg, Sarah S. Soliman, Matthew S. Kim, Devan H. Shah, Irene P. Chen, Melanie Ott, Wendell A. Lim, Hana El-Samad

## Abstract

Synthetic and chimeric receptors capable of recognizing and responding to user-defined antigens have enabled “smart” therapeutics based on engineered cells. These cell engineering tools depend on antigen sensors which are most often derived from antibodies. Advances in the *de novo* design of proteins have enabled the design of protein binders with the potential to target epitopes with unique properties and faster production timelines compared to antibodies. Building upon our previous work combining a *de novo*-designed minibinder of the Spike protein of SARS-CoV-2 with the synthetic receptor synNotch (SARSNotch), we investigated whether minibinders can be readily adapted to a diversity of cell engineering tools. We show that the Spike minibinder LCB1 easily generalizes to a next-generation proteolytic receptor SNIPR that performs similarly to our previously reported SARSNotch. LCB1-SNIPR successfully enables the detection of live SARS-CoV-2, an improvement over SARSNotch which can only detect cell-expressed Spike. To test the generalizability of minibinders to diverse applications, we tested LCB1 as an antigen sensor for a chimeric antigen receptor (CAR). LCB1-CAR enabled CD8+ T cells to cytotoxically target Spike-expressing cells. We further demonstrate that two other minibinders directed against the clinically relevant epidermal growth factor receptor are able to drive CAR-dependent cytotoxicity with efficacy similar to or better than an existing antibody-based CAR. Our findings suggest that minibinders represent a novel class of antigen sensors that have the potential to dramatically expand the sensing repertoire of cell engineering tools.

## Introduction

Synthetic receptors enable engineered cells to detect specific signals in their environment and respond with therapeutic payloads to treat disease^1–3^. Due to their therapeutic potential, synthetic biology groups have produced many different synthetic receptor systems. Perhaps the most widely exploited of these systems is the chimeric antigen receptor (CAR)^4^, which couples an antigen-sensing binder with signaling domains from the T cell receptor complex and associated co-stimulatory molecules. CAR T cells have proven tremendously valuable as anti-cancer therapeutics^5,6^. More recently, synthetic receptors such as synNotch^7^, MESA^8^, GEMS^9^, SNIPRs^10^, and many others^1^ were developed to use similar antigen-sensing domains coupled to customizable transcriptional programs. These synthetic receptor systems have shown great promise in enhancing existing cell-based therapies^11–16^ and bioengineering^2^ applications ranging from building cell-based biosensors^17–19^ to understanding and engineering development^20–26^.

Current receptors primarily rely on single chain variable fragment antibodies (scFvs) or less frequently variable heavy domain monomeric antibodies (nanobodies) to sense their targets. Monomeric antibodies that are generated for both research and therapeutic purposes are now mostly discovered through high-throughput screening using phage^27–29^ or yeast display^30^ followed by sub-screening in an orthogonal assay and then affinity maturation. Although these strategies have proven useful for generating antibody therapeutics and research tools, scFvs and nanobodies do not always translate easily to antigen sensors for synthetic receptors. While it is unclear why not all scFvs and nanobodies function in cell engineering contexts, it seems likely that the physical properties of binding, including affinity, on- and off-rates, and the steric and conformational requirements for each antibody, all have a role to play^31,32^. Further, although nanobodies and scFvs are relatively compact, even their modest size of >100 amino acids can be a limitation in the engineering of primary cells where genetic payload size is heavily constrained. For these reasons, small protein binders that can be engineered to bind specific targets hold promise as the next generation of antigen sensors.

*De novo*-designed binders that can be created using computational methods against almost any protein with known structure are a tantalizing new class of proteins that could serve as antigen sensors for synthetic receptors. Computational design of protein-protein interactions has a long history^33–35^ but recent advances have made this technology readily accessible for synthetic biology applications. Protein docking-based methods have enabled the design of binders against any protein structure^36^. Deep learning-assisted methods improved the speed of binder design and have yielded nanomolar affinity or better binders directly from computational pipelines without the need for affinity maturation^37–39^. Deep learning methods have also extended the design of binders to small helical peptides^40^, intrinsically disordered regions^41^dramatically expanding the scope of potential binder targets.

Previously, we demonstrated that a *de novo*-designed binder created using a docking-based approach that targeted the SARS-CoV-2 Spike protein^42^ could be combined with synNotch to drive customizable responses in a model of SARS-CoV-2 infection^18^. The SARS-CoV-2 minibinder used in those studies, LCB1, is a small three-helix bundle of 56 amino acids and binds directly to the interface it was designed for with picomolar affinity, overcoming many of the potential challenges for antigen sensor design. If minibinders like LCB1 generalize as antigen sensors across other synthetic receptors, these *de novo*-designed proteins could serve as powerful additions to the cell engineering toolbox.

In this study, we sought to test the generalizability of the LCB1 minibinder across different synthetic receptors. First, we demonstrate that LCB1 functions as an antigen sensor for a synthetic intermembrane proteolysis receptor (SNIPR)^10^ with distinct activation mechanics compared to synNotch. Unlike our previously described SARSNotch, the LCB1-SNIPR is capable of detecting both cells expressing the SARS-CoV-2 Spike and live SARS-CoV-2 virus. We also show that LCB1 functions as an antigen sensor for a CAR, demonstrating that this minibinder’s capabilities are not limited to proteolytic synthetic receptors but are also effective at modulating endogenous cellular pathways.

Minibinder-coupled CARs were able to specifically activate T cell programs in primary human CD8+ T cells against SARS-CoV-2 Spike-expressing cells. We further show the functionality of minibinders across targets and demonstrate the functionality of two anti-epidermal growth factor receptor (EGFR) CARs using minibinders targeted at two sites in the EGFR extracellular domain. Together, these findings suggest that minbinders can be easily deployed across synthetic receptors as antigen sensors.

## Results

### Minibinders adapt readily to next-generation synthetic proteolytic receptors

We previously demonstrated that the *de-novo* designed LCB1 and LCB3 minibinders for SARS-CoV-2 Spike protein could be used as functional antigen sensors when integrated with synNotch^18^. These anti-SARS Notch constructs successfully detected both purified Spike protein and Spike expressed on the surface of mammalian cells in a model of virally-infected cells to drive a synthetic transcriptional output.

We sought to extend these findings by testing the ability of the LCB1 and LCB3 minibinders to serve as antigen sensors for other synthetic receptors. Since the development of the original synNotch, a modular toolkit of parts for designing proteolysis-based receptors has been developed, allowing for tuning of receptor activation in various contexts^10^. These synthetic intramembrane proteolysis receptors (SNIPRs) use a collection of extracellular, transmembrane, juxtamembrane, and intracellular domains to tune baseline activity and receptor dynamic range for a variety of cellular and antigen contexts. Although these receptors function similarly to synNotch, the absence of the Notch regulatory repeats in the extracellular domain of SNIPRs suggests that their activation is mechanistically unique.

Given the similar function but mechanistic differences in activation of SNIPRs, we hypothesized that these receptors would be a good test case for evaluating minibinder function across synthetic receptors.

We selected a SNIPR chassis using the CD8a hinge domain, the human Notch1 transmembrane domain, and the human Notch2 juxtamembrane domain that had displayed low baseline activation and high payload expression upon specific activation in Jurkat and primary human CD4+ and CD8+ T cells. We generated LCB1- and LCB3-equipped SNIPRs coupled to Gal4-VP64 transcription factor (Figure 1A). We transduced these constructs into a Jurkat line expressing an output circuit as described previously^18^ that expresses BFP downstream of Gal4-VP64 activation. SNIPR-expressing

**Figure 1:**
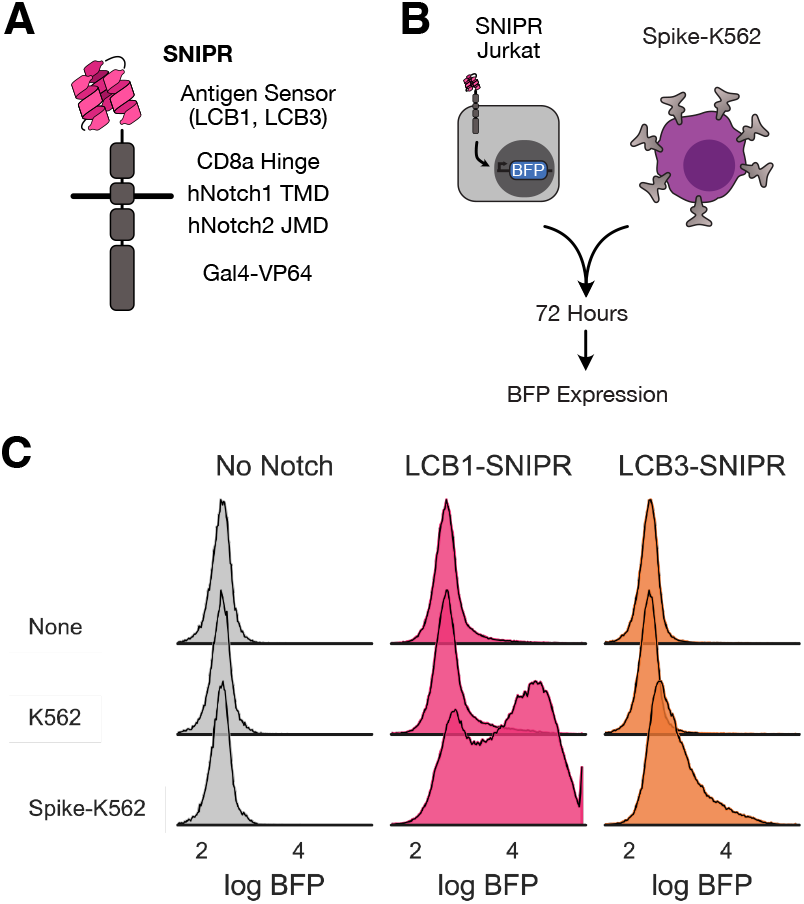
Minibinders adapt readily to next-generation synthetic proteolytic receptors. A) Schematic representation of a synthetic intramembrane proteolysis receptor (SNIPR) that enables customized genetic responses upon binding a target. LCB1 binder structure cartoon loosely based on PDB 7JZU. B) Assay for testing SNIPR function, whereby SNIPRs with various minibinder antigen sensors are expressed in Jurkat cells and cells are mixed with SARS-CoV-2 Spike-expressing K562 cells. SNIPR activation is readout as BFP expression. C) Histograms of BFP expression in SNIPR-expressing Jurkats after 72-hour incubation with K562s or K562s expressing Spike.

Jurkat T cells were sorted for surface staining of the expressed receptor, positive expression of a co-transduced mCherry marker, positive expression of mCitrine marking the integration of the output circuit, and the absence of BFP expression at baseline. See Supplemental Figure 1A for schema of output circuit and SNIPR expression constructs.

We incubated a polyclonal population of SNIPR-expressing cells with either untransduced K562 cells or K562s stably expressing a prefusion stabilized version of the SARS-CoV-2 Spike ectodomain^43^ displayed on a PDGFR transmembrane domain as previously described^18^ (Figure 1B). After 72 hours, we assessed our output circuit-only expressing Jurkat cells (No Notch) and LCB1- and LCB3-SNIPR expressing cells for BFP expression (Figure 1C). BFP expression was specifically increased for both SNIPR-expressing cell lines only in the presence of Spike-expressing K562s, demonstrating that minibinder-coupled SNIPRs specifically detected Spike protein on other cells.

### Minibinders perform similarly across different synthetic proteolytic receptors

Given the putatively different modes of activation between synNotch and SNIPRs, we wanted to compare the performance of minibinders between these different synthetic proteolytic receptors.

We sought to test performance in different antigen density regimes to better understand the dynamic range of the sensing capabilities of these minibinder-coupled receptors. We generated two Spike-expressing K562 populations expressing a moderate (Spike-K562 (M)) and high (Spike-K562 (H)) amount of Spike (Figure 2A). We also regenerated the LCB1-Notch and LCB3-Notch expressing Jurkat lines described in our previous work to match the expression level of LCB1- and LCB3-SNIPR in the lines generated above (Figure 2B, Supplemental Figure 1A). We then compared polyclonal populations of these lines in the same 72 hour incubation assay described above.

**Figure 2:**
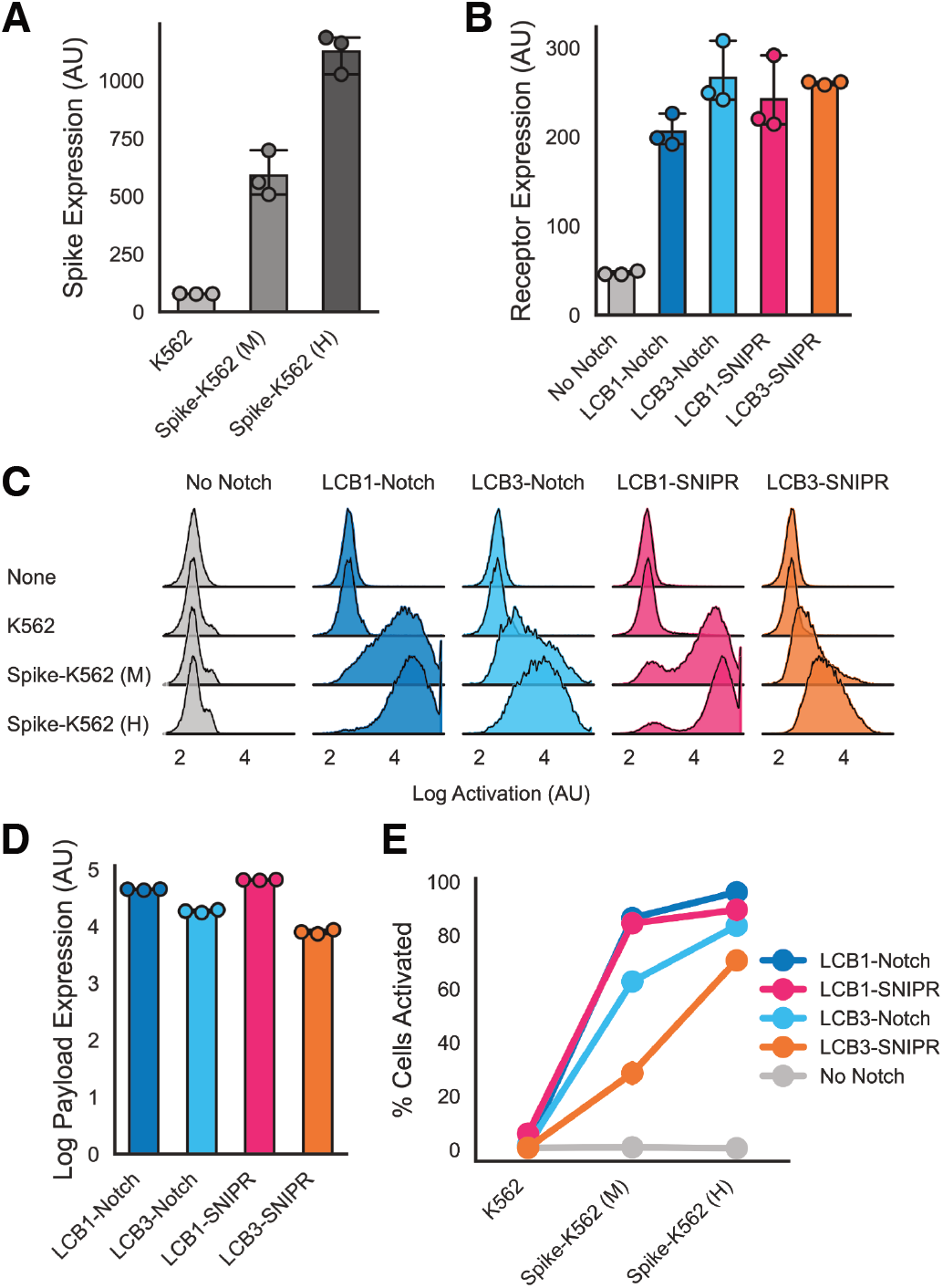
Minibinders perform similarly across different synthetic proteolytic receptors. A) Expression of surface-displayed Spike protein (as assessed by anti-Myc labeling of an N-terminal extracellular Myc tag) for untransduced K562s and K562 lines sorted for moderate (M) and high (H) expression of Spike (median fluorescent counts for populations across 3 replicates). B) Surface expression levels of multiple minibinder-coupled synthetic receptors on Jurkat cells as assessed by anti-Myc labeling of an N-terminal extracellular Myc tag on each receptor (median fluorescent counts for populations across 3 replicates). C) Histograms describing BFP expression in minibinder receptor-expressing Jurkats after 72 hours of incubation with K562s with 3 different levels of Spike expression (pooled populations from 3 replicates). D) Mean log BFP expression in the activated populations of Jurkats after 72 hour incubation with Spike-K562 (H) cells as calculated by the estimated mean of the high BFP expression population after fitting a two component Gaussian mixture model on each of 3 replicates per condition. Data points are estimated mean of ‘on’ population in each replicate. E) The fraction of BFP-expressing Jurkats in the mixture model-defined activated population for each receptor as a function of Spike expression. All error bars (including E) are standard deviation.

All minibinder-coupled receptor lines expressed BFP specifically in response to both M and H Spike-expressing K562s (Figure 2C). To differentiate activated (BFP-expressing cells) from inactive cells, we fit a two component Gaussian mixture model to the BFP expression data across experimental populations. We compared the predicted mean of the activated component identified by the mixture model across each receptor to understand the amount of payload that each receptor could produce in response to Spike-K562s (Figure 2D). For both synNotch and SNIPR, LCB1-coupled receptors produced greater payload expression in response to Spike-expressing cells compared to LCB3-coupled receptors. SynNotch and SNIPR receptors coupled to the same minibinder produced similar levels of BFP expression. To determine the fraction of each population successfully activated (expressing BFP), we fit our mixture model to each experiment and assessed the fraction of cells in the population with the higher mean expression (Figure 2E). All receptors showed an antigen dose-dependent increase in activation. Again, LCB1-coupled receptors showed the highest % activation, with LCB1-SNIPR (90% ± 0.5% activation against Spike-K562 (H) cells) and LCB1-Notch (96% ± 0.3%) performing similarly, followed by LCB3-Notch (84% ± 0.5%) and then LCB3-SNIPR (71% ± 1.9%) with the lowest activation.

These findings suggest that despite distinct mechanisms of activation of synNotch and SNIPRs, minibinders perform similarly across these receptors. This is true both for the maximum efficacy of a specific minibinder, and also the relative performance of two different minibinders directed at the same target.

### Novel synthetic proteolytic receptors enable minibinder-dependent detection of live SARS-CoV-2 virus

Although minibinders performed similarly between synNotch and SNIPRs in a cell-expressed antigen context, we hypothesized that the distinct activation mechanism of SNIPRs might provide advantages for detection in other contexts. Previously, we showed that LCB1-Notch was not capable of activating in response to pseudotyped lentivirus decorated with the SARS-CoV-2 Spike. We sought to determine whether SNIPR’s unique activation mechanism would enable it to better detect SARS-CoV-2 virus itself.

Using the Jurkat lines generated above, we took our best-performing receptors LCB1-Notch and LCB1-SNIPR and incubated them with live SARS-CoV-2 virus for 72 hours before fixing and assessing BFP expression (Figure 3A). We initially incubated receptor-expressing Jurkats with no virus, the WA1 viral isolate, or the B.1.617.2 (Delta) virus isolate, all at a multiplicity of infection of 1. When assessing median BFP expression (Figure 3B) we saw that LCB1-Notch produced no BFP in response to virus, LCB1-SNIPR had higher baseline activity, but importantly it also showed increased BFP expression in response to both viral isolates. We then conducted a dose-response study, incubating LCB1-SNIPR cells with a 2 log range of viral MOIs and assessing the fraction of cells that activated (Figure 3C). LCB1-SNIPR cells showed a modest dose-dependent increase in % cells activated for both viral isolates, with stronger activation for WA1.

**Figure 3:**
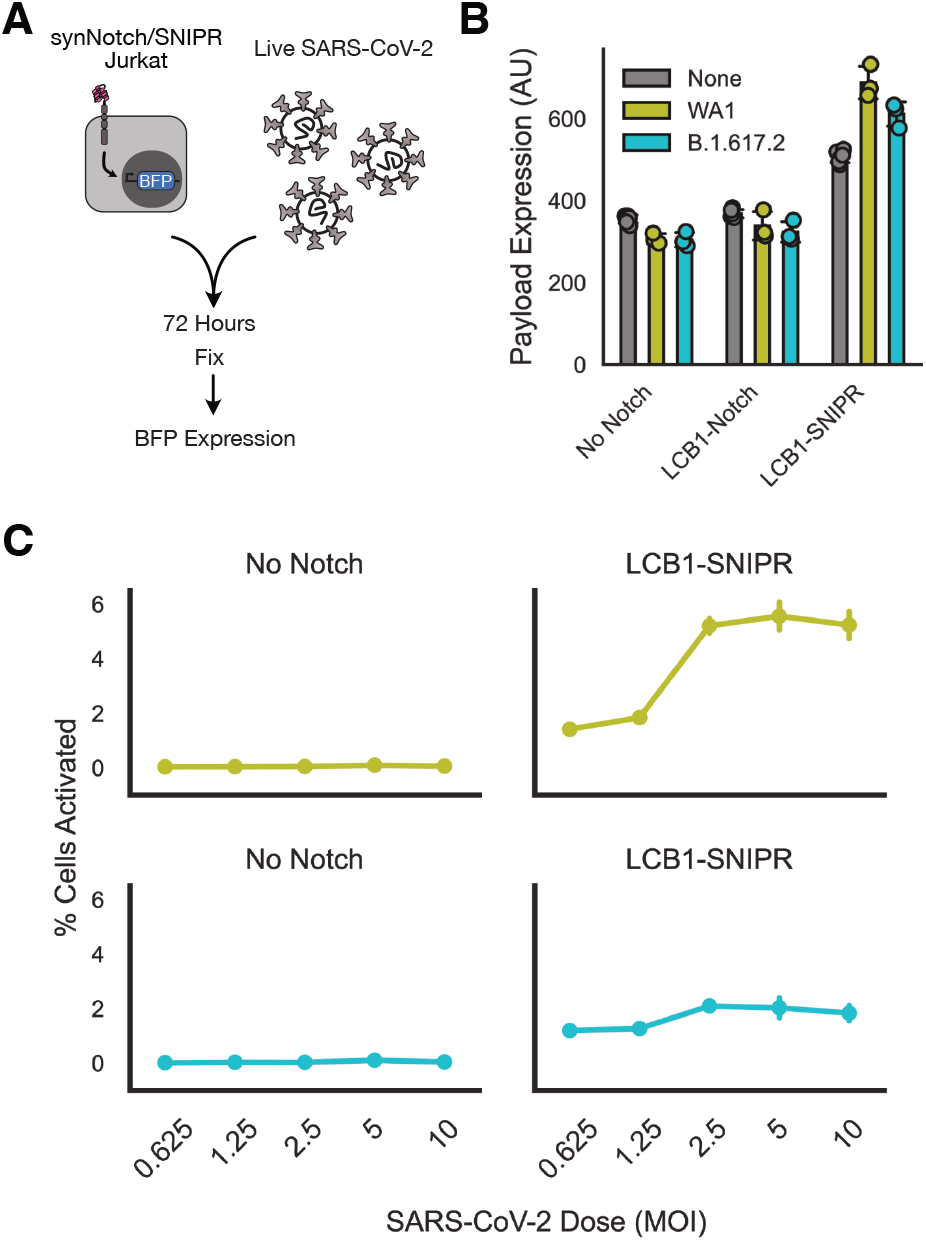
Novel synthetic proteolytic receptors enable minibinder-dependent detection of live SARS-CoV-2 virus. A) Schematic of live virus sensing experiment. Minibinder receptor-expressing Jurkats were incubated with live SARS-CoV-2 virus for 72 hours, and then fixed with paraformaldehyde and assessed for BFP expression as a sign of receptor activation. B) Median BFP expression in Jurkat cells exposed to no virus or WA1 or Delta isolates of SARS-CoV-2 at a multiplicity of infection of 1. C) Fraction of Jurkat population expressing BFP as a function of MOI of SARS-CoV-2 that cells were exposed to. All error bars are standard deviation.

These data suggest that although performance of these minibinders generalizes well across synthetic receptors, the minibinders can successfully enable unique capabilities for the different receptors they are coupled to.

### Minibinders enable chimeric antigen receptor-mediated killing of SARS-CoV-2 Spike-expressing cells

After demonstrating minibinder generalization to multiple proteolytic receptors, we sought to extend these findings to a distinct receptor class to test minibinder viability as antigen sensors across a wider range of synthetic receptors. For this study, we selected the chimeric antigen receptor (CAR) because of its important therapeutic relevance.

We generated a minibinder-coupled second-generation CAR by combining LCB1 with the CD8α hinge and transmembrane domains and the intracellular domains of 4-1BB and CD3ζ^44^ (Figure 4A). This construct, dubbed LCB1-CAR, expressed well when lentivirally transduced into CD8+ primary human T cells (Supplemental Figure 2B).

**Figure 4:**
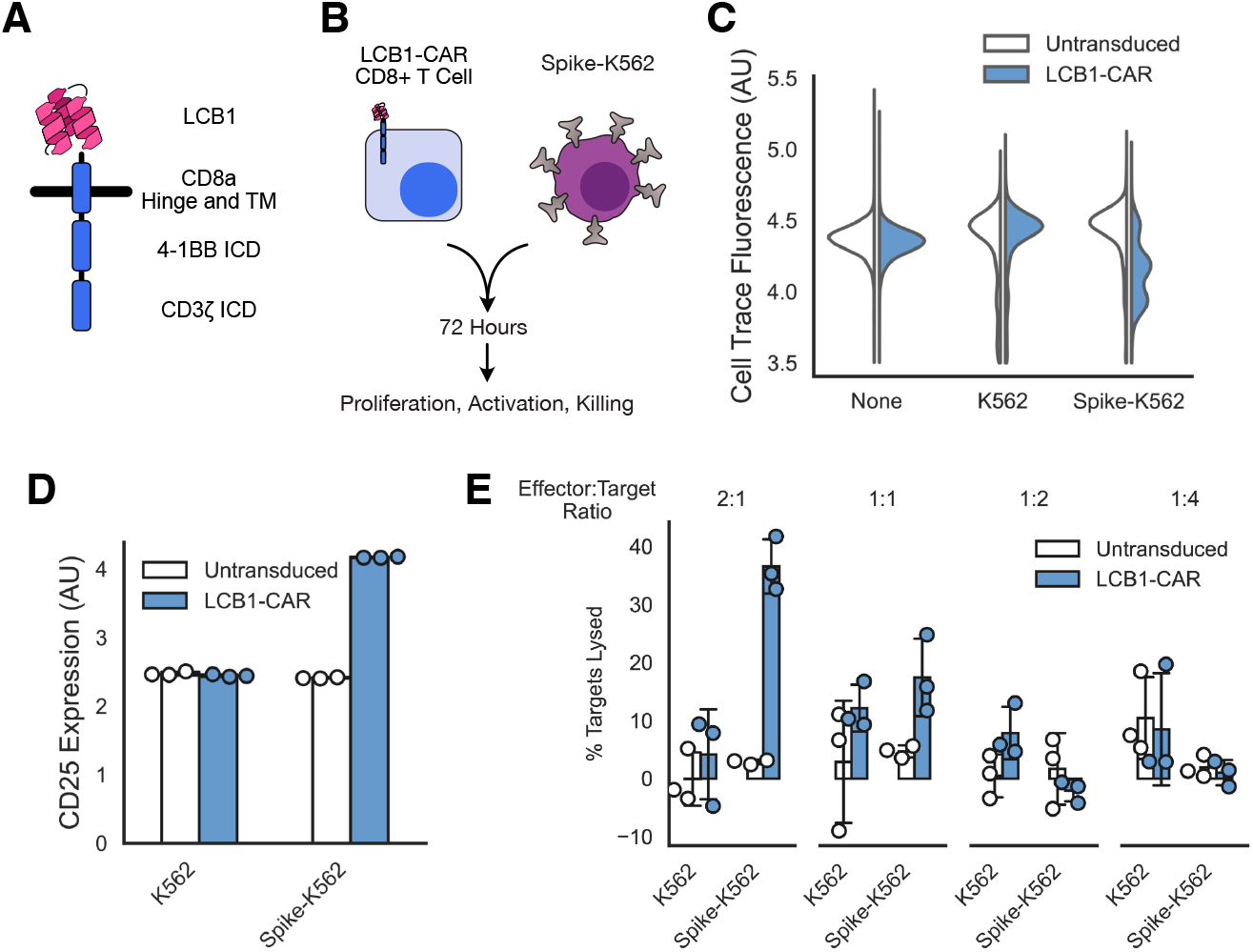
Minibinders enable chimeric antigen receptor-mediated killing of SARS-CoV-2 Spike-expressing cells. A) Schematic of a chimeric antigen receptor with 4-1BB costimulatory domain using the minibinder LCB1 as its antigen sensor. B) Assay schematic indicating that primary CD8+ human T cells expressing the LCB1-CAR were mixed with Spike-expressing K562s for 72 hours and then were assessed for proliferation, T cell activation, and killing of the target cells. C) Proliferation of T cells as a function of which target cells they were incubated with. Histogram showing Cell Trace dye fluorescence across a population of T cells, with homogenous high staining indicating no proliferation and multiple populations indicating proliferation. D) T cell activation as assessed by CD25 antibody-labeling as a function of which targets T cells were exposed to (median fluorescent for population in each of 3 replicates). E) The fraction of target cell populations that were lysed after T cell incubation as a function of the ratio of effector T cells to K562 targets that were incubated together. All error bars are standard deviation.

We tested the ability of LCB1-CAR to induce an antigen-specific cytotoxic phenotype (Figure 4B). After transducing CD8+ T cells with LCB1-CAR, we co-cultured them with either parental K562 cells or Spike-K562 (H) cells. We co-cultured our effector and target cells at ratios ranging from 2 effectors for every target to 4 targets for every effector. We then evaluated the ability of LCB1-CAR to elicit cell proliferation, expression of the T cell activation marker CD25, and lysis of target cells.

LCB1-CAR promoted a specific cytotoxic reaction to Spike-expressing K562s (Figure 4C-E). LCB1-CAR-expressing T cells proliferated in the presence of Spike-K562s but not when cultured on their own or with parental K562s, as indicated by a dilution of the CellTrace labeling dye in cell populations only in the LCB1-CAR/ Spike-K562 condition (Figure 4C). The expression of the activation marker CD25 was upregulated for LCB1-CAR cells in the presence of Spike-K562s but not in other conditions (Figure 4D). Importantly, LCB1-CAR T cells were able to induce specific lysis of Spike-K562s at effector:target ratios of 2:1 (∼40% lysis) and 1:1 (∼20% lysis), with little off-target lysis of other cell types (Figure 4E). These data together demonstrate that the minibinder LCB1 is a functional antigen sensor for a CAR, albeit with moderate efficacy.

### Short linker sequences optimize minibinder-coupled receptor function

We next sought to improve the efficacy of minibinder-coupled CARs. Although LCB1 proved the better of the two binders in proteolytic receptor function, we considered that LCB3 might have different efficacy when coupled to a CAR. LCB1 and LCB3 bind the same interface on the Spike protein, but in opposite orientations^42^. We hypothesized that this binding difference, which was detrimental to LCB3 function on synNotch and SNIPR, might provide different activity for a CAR.

In comparing LCB1- and LCB3-CARs, we also added in a CD19-CAR as a benchmark, as this CAR is the gold standard in the field. Our CD19-CAR used the same 4-1BB and CD3ζ intracellular domains as the LCB-CARs, swapping in the CD19 scFv FMC63^45^ for the antigen sensor domain. All of the LCB1-, LCB3-, and CD19-CARs expressed at similar levels after lentiviral transduction into primary human CD8+ T cells (Supplemental Figure 2B).

We evaluated cytotoxicity of each of these CARs against K562s, CD19-K562s (Supplemental Figure 2A), and Spike-K562 (H) cells (Figure 5A). As expected, the CD19-CAR performed well and lysed 96% ± 0.5% of target cells. Both LCB-CARs performed similarly to each other and at about half the efficacy of the CD19-CAR, with LCB1-CAR lysing 43% ± 6.4% of targets and LCB3-CAR lysing 38% ± 6.9% of targets. Efficacy varied between T cells from different donors, with all 3 CARs showing similar specific cytotoxicity when expressed in donor cells that had higher levels of off-target cytotoxicity (Supplemental Figure 2C). Together, these results suggest that unlike with synNotch and SNIPR, the binding orientation of LCB1 and LCB3 do not affect CAR function, and that these minibinders are less efficacious antigen sensors compared to the gold-standard CD19 scFv.

**Figure 5:**
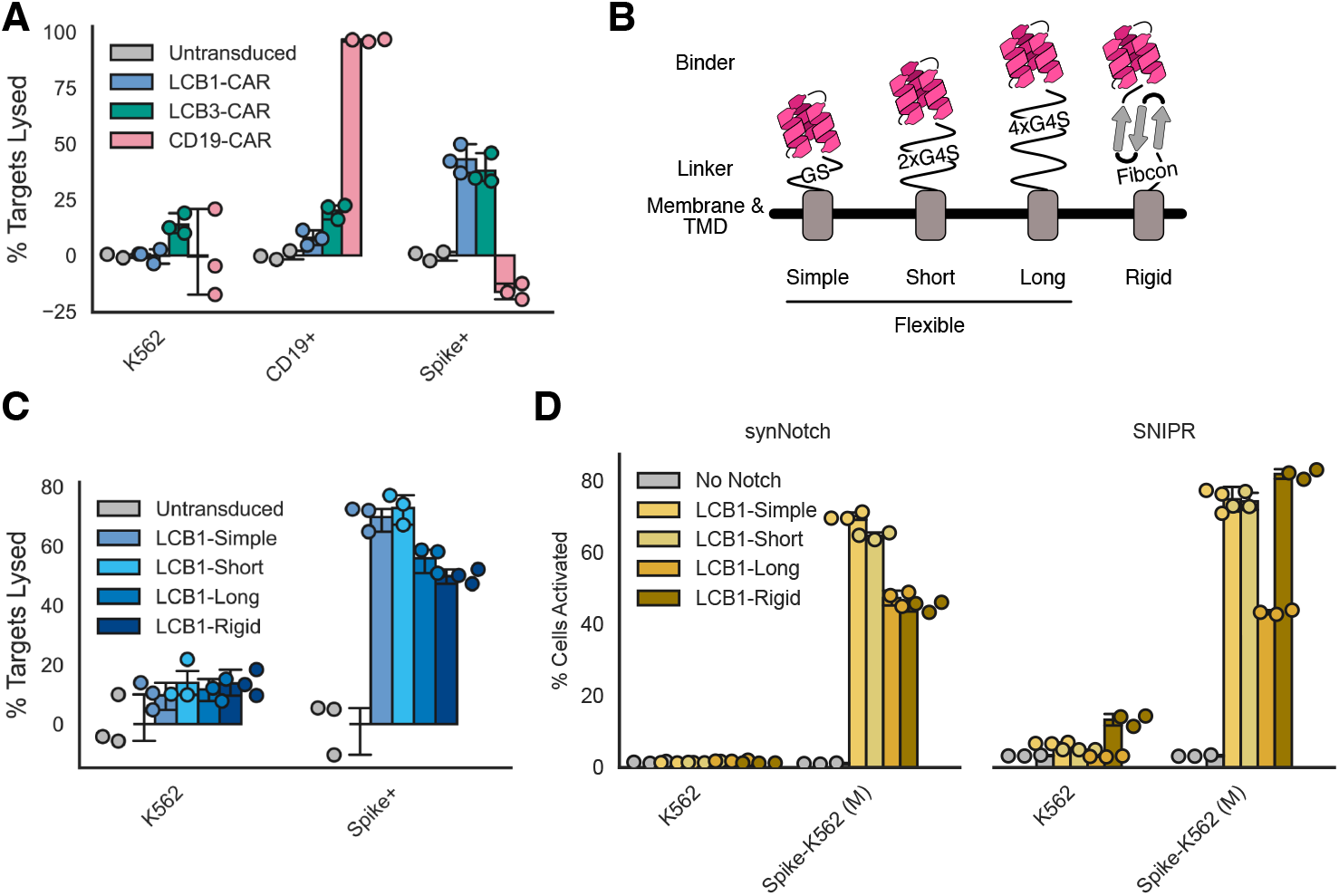
Short linker sequences optimize minibinder-coupled receptor function. A) The fraction of K562s expressing no ligand, CD19, or Spike that were lysed after 72 hour incubation with LCB1-CAR, LCB3-CAR, or the benchmark CD19-CAR at a 1:1 effector:target ratio. B) Schematic describing 4 different linkers that were evaluated to explore the effect of linker length and rigidity on minibinder receptor function. Fibcon linker structure cartoon loosely based on PDB 3TEU. C) The fraction of K562 of Spike-K562 targets lysed after 24 hour incubation with CD8+ T cells expressing 4 variants of the LCB1-CAR at a 1:1 effector:target ratio. D) The percent of proteolytic receptor-expressing cells that were expressing BFP after 72 hour incubation with K562 or Spike-K562 (M) target cells for 4 different linker variations. All error bars are standard deviation.

We next sought to test optimizations in other aspects of the binders. Antigen sensor affinity^46,47^ and the composition of the linker region between the antigen sensor and the CD8a hinge^46,48^ have both been demonstrated to increase CAR-dependent lysis of target cells. Both LCB minibinders have picomolar affinity for their targets, similar to or better than the ∼300pM affinity described for the CD19 scFv we used^49^ which suggests increasing affinity is unlikely to improve performance.

Instead, we targeted the linker region (Figure 5B). Our initial construct featured a minimal linker (GS, “simple”) between the minibinder and the CD8a hinge domain used in the CAR. We generated constructs including a short flexible linker (2x repeats of GGGGS, “short”), a longer flexible linker (4x repeats of GGGGS, “long”), and a synthetic stabilized immunoglobulin fold-like domain^50^ predicted to be highly stable as a rigid alternative to the flexible linkers above (Fibcon, “rigid”). We constructed LCB1-based CARs with each of these linkers, and they all expressed at similar levels (Supplemental Figure 2B). We then evaluated these minibinder and linker variants in the same killing assay described above (Figure 5C). At hours after co-culture with target cells, mild differences could be seen between linker variants, with modestly lower cytotoxicity from the long and rigid variants compared to the simple and short variants. By 72 hours post co-culture, these differences disappeared and all variants behaved similarly (Supplemental Figure 2D). Altogether, the simple GS linker appears to be optimal for CAR-dependent cytotoxicity.

With these linker variants in hand, we also explored whether these linkers would affect minibinder-coupled synNotch or SNIPR function. All LCB1 linker variants successfully expressed when coupled to synNotch or SNIPR, but lower expression was seen for the short, long, and rigid variants when coupled to SNIPR (Supplemental Figure 3A). We then tested Jurkats expressing these variants in a co-culture experiment with Spike-K562 (M) cells (Figure 5D). Expression differences did not correlate with differences in functionality. LCB1-Simple and -Short versions of synNotch functioned best, while the LCB1-Long and -Rigid activated a much smaller percentage of cells in response to Spike-K562s (∼75% of cells activated for LCB1-Simple and -Short Notch vs ∼45% activated for LCB1-Long and -Rigid Notch). SNIPR was similarly unable to tolerate the long linker, but LCB1-Rigid SNIPR performed similarly to the variants with the simple and short linkers.

These data suggest that while minbidiners function well across a variety of synthetic receptors, there is still more exploration needed to determine how to create minibinders with the best efficacy at chimeric antigen receptors. Overall, our data suggest that minibinders work best with a short flexible linker coupling them to the receptor they are serving as antigen sensors for.

### EGFR minibinders enable CAR-mediated recognition of a clinically relevant target

We next explored whether minibinders could drive receptor function across different antigen targets. De novo-deisgned minibinders have been produced for over a dozen different naturally occuring proteins^36,39,40^. From these existing minibinders, we selected two binders to evaluate that were designed against the epidermal growth factor receptor (EGFR), a clinically relevant target in cancer^51^ that has been previously targeted using CARs.^47,52,53^. The two binders, EGFRn_mb and EGFRc_ mb^36^, were designed against the N-terminal and C-terminal structured domains of EGFR’s extracellular face (Figure 6A).

**Figure 6:**
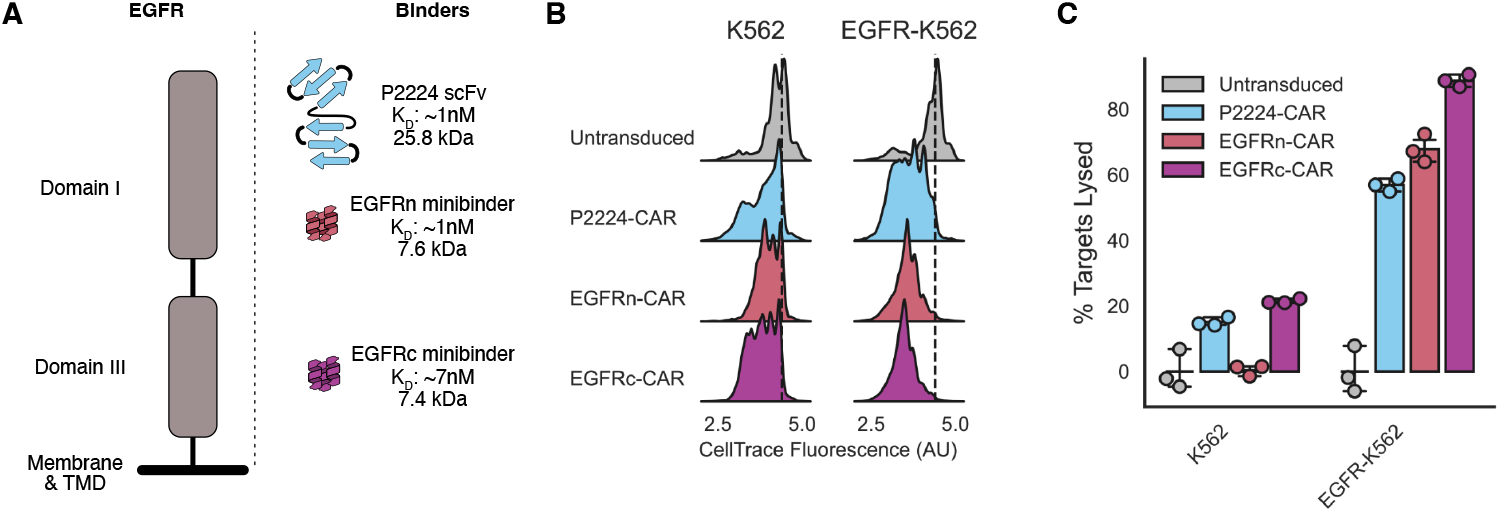
EGFR minibinders enable CAR-mediated recognition of a clinically relevant target. A) Schematic representation fo the EGFR extracellulaor domain and the binders used in these experiments. Each of the binders is placed according to its predicted binding site on EGFR. B) Proliferation of T cells as a function of which CAR they are expressing and which target cells they were incubated with. Histogram showing Cell Trace dye fluorescence across a population of T cells, with homogenous high staining indicating no proliferation and multiple populations indicating proliferation. Dotted line demotes location of unproliferating population. C) The fraction of target cell populations that were lysed after T cell incubation as a function of which cell type T cells were incubated with. All error bars are standard deviation.

To accurately assess the efficacy of these EGFR minibinders as CAR antigen sensors, we compared them to an existing anti-EGFR CAR. We selected a CAR coupled to the affinity-matured P2224 scFv^54^ which has been previously shown to induce lysis of a variety of EGFR-expressing cells lines when coupled to a CAR^47^.

The features of the binders including their size, affinity, and predicted location of binding are schematized in Figure 6A. As noted previously, the minibinders are roughly ⅓ the size of the scFv. All binders have similar measured affinities to their target at <10nM, albeit assessed by different methods (binding of fluorescently labeled protein to EGFR-expressing A431 cells for P2224^54^, biolayer interferometry for the minibinders^36^). Each of the binders are predicted to bind to distinct regions of the EGFR extracellular domain^55^: P2224 to the distal N terminus, EGFRn to the middle of structured Domain I, and EGFRc to the middle of structured Domain III.

We generated CARs with each of the minibinders and then expressed them or the P2224-CAR in primary human CD8+ T cells to conduct the same cytotoxicity assay as described above for the LCB CARs. Both minibinder CARs had improved surface expression compared to the P2224 CAR (Supplemental Figure 4A). We tested the ability of the CARs to mediate T cell activation in the presence of either untransduced K562s or K562s transduced to express the EGFR extracelluar domain displayed on a PDGFR transmemberane domain (Supplemental Figure 2B). Effector T cells and target K562s were co-cultured as described above for 72 hours and then assessed for proliferation and target cell lysis. All CAR-expressing cells showed modest proliferation in the presence of untransduced K562s compared to untransduced T cells, while all showed further specific proliferation when incubated with EGFR-K562s (Figure 6B). The P2224- and EGFRc-CARs showed modest off-target killing of K562s, while all 3 CARs showed similar efficacy in specific lysis of EGFR-K562s (Figure 6B).

When assessed at different effector-to-target ratios where the number of target cells was left constant and the number of effector cells reduced across ratios, all CARs showed similar efficacy at each ratio with modestly better performance of the minibinder CARs at the lowest ratio of 1 effector to 16 targets (Supplemental Figure 4C). Overall, our data suggest that both EGFR minibinders perform similarly or better than an existing scFv CAR in our *in vitro* assays, futher demonstrating the value of minibinders as CAR antigen sensors.

## Discussion

In this study, we demonstrated that *de novo*-designed protein minibinders generalize as antigen sensors across multiple synthetic receptors. The LCB1 and LCB3 binders targeting SARS-CoV-2 Spike successfully induced activation of two proteolytic receptors in the presence of cells expressing Spike, and LCB1 enabled SNIPR-based detection of live SARS-CoV-2 virus. LCB1 and LCB3 similarly enabled chimeric antigen receptors expressed in human CD8+ T cells to recognize and drive killing of Spike-expressing cells in culture. We evaluated several potential optimizations to improve the performance of minibinder-coupled receptors and found that for the receptors tested short flexible linkers produced the optimal functionality. Finally, we demonstrated that two minibinders against EGFR enabled CAR-dependent cytoxicity at levels similar to or better than an existing scFv-based anti-EGFR CAR. Together, these results identify *de novo*-designed minibinders as an important part of the burgeoning cell engineering toolbox.

The two anti-Spike minibinders LCB1 and LCB3 showed differential efficacy across receptors. For both of the proteolytic receptors, LCB1 was a dramatically better antigen sensor and resulted in as much as 10-fold higher payload expression and twice as much population activity compared to the LCB3-coupled receptors. We previously hypothesized that this difference was the result of the binding orientation of these two proteins, which bind to the same surface on Spike but in opposite orientations^42^. Curiously, this hierarchy in function is not dependent on the activation mode of the proteolytic receptor – synNotch activation is thought to depend on its Notch-derive extracellular domain^7^ while the SNIPRs are hypothesized to function through dimerization and an as-yet unelucidated mechanism that is dependent on the CD8a extracellular hinge domain^10^. The shared functional hierarchy of LCB1 and LCB3 between these receptors suggests that despite their distinct mechanisms of activation, they share some common requirements of their antigen sensors. This hierarchy was not seen for the same binders on a CAR, where both LCB1 and LCB3 produced similar T cell activation profiles. Further insight is provided by the EGFR minibinders, which bind in similar orientations but to different parts of the EGFR ectodomain. As CAR antigen sensors, the EGFRc minibinder which binds to EGFR closer to its transmembrane domain, performed modestly better than the EGFRn minibinder despite both having similar affinity for their target. Our results suggest that a systematic exploration into the role of binding orientation might aid in creating optimal *de novo*-designed antigen sensors in the future.

Of the linkers we explored, our data suggest that the optimal linker sequence for connecting minibinders to a target receptor is a simple glycine-serine (GS) linker. All receptors tested tolerated both a GS linker and 2xGGGGS linker well, but the performance of a longer flexible linker was poor across all receptors, and only SNIPR tolerated a more rigid linker. The linkers tested in this study are not exhaustive – future studies incorporating minibinders into synthetic receptors could also explore distance from the plasma membrane as a key variable of linker length.

Although this is difficult to quantify in the context of this study, we want to note that all the minibinder-coupled receptors used here were rapidly developed. Each receptor included in this study went from protein sequence to functional assay in one month or less. From discussions with colleagues, we note that this is abnormal for generating proteolytic or chimeric antigen receptors against novel targets, where significant troubleshooting of both antigen sensor and linker length are often required. Indeed, in our own development of the original SARSNotch, several months were spent attempting to generate receptors using alternative scFv- or nanobody-based anti-Spike binders before the LCB1-Notch worked on first attempt. All the minibinders used in this study were previously biophysically characterized, but we suspect that novel methods for in-cell screening of minibinders with functional outputs hold considerable potential for rapidly developing novel minibinder-coupled receptors.

Computational approaches for creating *de novo-* designed binders against practically any protein^34,36–39,41,56– 59^ or peptide^40^ with known structure position minibinders as attractive new antigen sensors. The ability of these methods to target specific interfaces of proteins of interest will further enable novel synthetic receptors that can distinguish between different faces and perhaps even conformations of membrane-displayed proteins. Further, novel methods for designing high-affinity binders for small proteins like neuropeptides and peptide hormones^40^ have the potential to build synthetic receptors capable of robust activation in the presence of secreted factors, a sorely lacking piece of the current cell engineering toolkit. Further, although binders have only been designed against proteins with known structure, structure prediction methods^55,60,61^ might enable the design of binders against proteins without experimentally elucidated structures. *De novo*-designed binders are already seeing use both as pharmacological tools for controlling cell function^58^ as well as biosensors for measuring the activity of endogenous proteins^62^. We anticipate that as nanobodies have enabled intracellular sensing of proteins^63,64^ and their conformations^65–68^, *de novo*-designed binders will enable a new generation of intracellular tools that can control the stability^69–71^ and function^72–74^ of proteins intracellularly.

Together, we hope the minibinder-coupled receptors described here prompt the rapid adoption of novel minibinders against a diversity of targets to dramatically expand the perceptual repertoire of cell engineering tools.

## Materials and Methods

### DNA Constructs

The synthetic receptors and minibinders used in this study (described in Supplemental Table 1 and schematized in Supplemental Figure 1) were constructed using the Mammalian Toolkit (MTK)^75^, a hierarchical DNA assembly method. The SNIPR used in this study was a generous gift from Dr. Kole Roybal and contains the CD8a hinge domain, the human Notch1 transmembrane domain, and the human Notch2 juxtamembrane domain. The SNIPR was domesticated as an MTK part 3B with the Gal4-VP64 transcriptional actuator domain. The chimeric antigen receptor used in this study is comprised of the CD8a hinge and transmembrane domains, the 4-1BB costimulatory domain, and the CD3ζ chain and was also domesticated as an MTK part 3B. The anti-Spike minibinders were reused from modular parts plasmids previously described^18^, while the EGFR minibinders were generated by codon optimizing reported protein sequences^36^ and appending the same CD8a signal sequence and Myc epitope tag used previously on the N terminus. LCB1 linker variants were generated *de novo* as gene blocks ordered from IDT. The rigid linker is the Fibcon domain that has been previously described^50^ with a sequence of MLDAPTDLQVTNVTDTSITVSWTPPSATITGYRITYTPSNG PGEPKELTVPPSSTSVTITGLTPGVEYVVSVYALKDNQESP PLVGTQTTG. The anti-CD19 CAR was a generous gift from Dr. Andrew Ng. The EGFR extracellular domain displayed on PDGFR was a generous gift from Dr. Greg Allen. The anti-EGFR P2224 scFv CAR was a generous gift from Dr. Ki Kim.

For lentiviral transduction, a new set of MTK plasmids were generated as part of this study. The lentiviral plasmids previously used for SARSNotch^18^ were mutated to remove BsaI and BsmBI restriction sites. In the process of this study, we discovered that these plasmids were suboptimal for transfecting primary T cells. A novel MTK lentiviral backbone, pMK1101, was generated based on the pHR series of lentiviral plasmids that used the PaqCI restriction sites for assembly. A new series of transcriptional unit vectors (pMK1213, pMK1129, pMK1130, pMK1131, pMK1132) were generated that allow transcriptional unit assembly via the standard BsaI MTK Golden Gate reaction that can then be assembled into pMK1101 via a PaqCI reaction following manufacturer instructions (New England Biolabs #R0745S).

All plasmids were propagated in Stbl3 *E. coli* (QB3 MacroLab). Domestication was verified via sequencing and transcriptional unit assembly was verified via restriction digest and sequencing. All plasmids will be available on Addgene, and the sequence for all constructs used in this study are included as supplemental data.

### Cell Culture

K562 and Jurkat cell lines were cultured in RPMI-1640 media (Gibco #11875-093) supplemented with 10% fetal bovine serum (FBS) (Hyclone #SH30071.03), and 1% Anti-Anti (Gibco #15240-096).

Primary human CD8+ T cells were isolated from anonymous donor blood after apheresis by negative selection. T cells were cryopreserved in RPMI-1640 (Corning #10-040-CV) with 20% human AB serum (Valley Biomedical, #HP1022) and 5% DMSO (Sigma-Aldrich #472301). After thawing, T cells were cultured in human T cell medium consisting of X-VIVO 15 (Lonza #04-418Q), 5% Human AB serum and 10 mM neutralized N-acetyl L-Cysteine (Sigma-Aldrich #A9165) supplemented with 30 units/mL IL-2 (NCI BRB Pre-clinical Repository).

Lenti-X 293T packaging cells (Clontech #11131D) were cultured in medium consisting of Dulbecco’s Modified Eagle Medium (DMEM) (Gibco #10569-010) and 10% fetal bovine serum (FBS) (University of California, San Francisco [UCSF] Cell Culture Facility). Lenti-X 293T cells were cultured in 150 or 225 flasks (Corning #430825 and #431082) and passaged upon reaching 80% confluency. To passage, cells were treated with TrypLE express (Gloco #12605010) at 37 C tor 5 minutes. Then 10 mL of media was used to quench the reaction and cells were collected into a 50 mL conical tube and pelleted by centrifugation (400xg for 5 minutes). Cells were cultured until passage 30 whereupon fresh Lenti-X 293T cells were thawed.

Vero-TMPRSS2 obtained from ATCC were cultured in DMEM (Corning) supplemented with 10% fetal bovine serum (FBS) (GeminiBio), 1% glutamine (Corning), and 1% penicillin-streptomycin (Corning).

All cells were maintained at 37°C with 5% CO2, and cell counting was conducted using a Trypan Blue (Invitrogen #T10282) exclusion stain and an automated Countess II FL (Invitrogen #AMQAF1000) cell counter.

### Lentiviral Transduction

Lenti-X 293T cells (Takara Bio #632180) were seeded at approximately 7e5 cells/well in a 6-well plate to yield ∼80% confluency the following day. The following day cells were transfected with 1.5μg of transfer vector containing the desired expression cassette, and the lentiviral packaging plasmids pMD2.G (170ng) and pCMV-dR8.91(1.33μg) using 10μl of Fugene HD (Promega #E2312) according to manufacturer protocols. At 48 hours the viral supernatant was filtered through a 0.45μm PVDF filter and added to Jurkat or K562 cells seeded at approximately 1e5 cells/well in a 12-well plate. After 24 hours, the viral media was then exchanged for fresh media. Polyclonal cell populations were selected via fluorescence activated cell sorting (FACS) as described below.

For each construct tested, 1e6 Primary T cells were thawed and activated the next day using CD3/CD28 Dynabeads (Life Technologies #11131D) at a 1:3 cell: bead ratio. Lenti-X 293T cells were transfected to generate virus the same day that T cells were activated. At 48 hours post-transfection, the viral supernatant was filtered through a 0.45μm PVDF filter and concentrated using the Lenti-X concentrator (Takara #631231) according to the manufacturer’s instructions. For each construct, 2 wells of a 6-well plate of virus were concentrated together and then incubated in one well of a 24-well plate with 1e6 T cells. 24 hours after the addition of virus to T cell culture, viral media was removed and fresh media was added. At day 5 post T cell stimulation, Dynabeads were removed and the T cells were sorted for construct expression via FACS. T cells were then expanded for at least eight days until assays.

### Antibodies

Surface-expressed proteins were assayed using Alexa Fluor 647 Anti-Myc tag antibody (Cell Signaling Technologies #2233S). T cell activation was assessed using Alexa Fluor 647 anti-CD25 (Biologend #302618). CD19 expression was assessed using FITC anti-CD19 (Biolegend #302206). EGFR extracellular domain expression was assessed using BV786 anti-EGFR (BD Biosciences #749750). All antibodies were diluted 1:100 in DPBS (UCSF Cell Culture Facility) for staining.

### Fluorescence Activated Cell Sorting (FACS)

Cell lines were bulk sorted for high expression using the UCSF Laboratory for Cell Analysis Core Facility FACSAriaII (BD Biosciences) or a Sony SH800.

Fluorescence-negative controls were used to set detector power so that negative cells appeared to have mean fluorescence ∼100 counts, and then transduced cells were sorted for cells with expression outside of the negative control expression level. Jurkat cells transduced with synNotch and SNIPR output circuit were sorted for low BFP and high mCitrine expression. SynNotch and SNIPR transduced Jurkats cells were stained for surface expression and then sorted for low BFP, high mCherry, high surface expression, and high mCitrine expression.

Spike-expressing K562s were stained for surface expression and sorted for high surface expression and high mCherry. T cells transduced with CAR variants were stained for surface expression and then sorted for high surface expression and high mCherry.

### Flow Cytometry

Flow cytometry was performed using a LSR-Fortessa (BD Biosciences). Prior to the flow cytometry, cells were seeded at densities described below in 96 well U bottom plates (Falcon #877217) and incubated for 24-72 hours as specified by the experiment. Plates were then spun at 400xg for 5 minutes to settle all cells. Cells were then resuspended and stained for 45 minutes with appropriate antibodies. After incubation, plates were spun down, rinsed with 200µl DPBS, and resuspended in flow buffer (DPBS with 2% FBS). The plates were then run on the flow cytometer using a four laser configuration (405nm, 488nm, 561nm, 640nm), collecting fluorescence for TagBFP (405nm ex, 450/50 em), mCitrine (488nm em, 530/30 em, 505lp dichroic), mCherry (561nm ex, 610/20 em, 600lp dichroic), and Alexa Fluor 647 (640nm ex, 670/14 em). At least 10,000 events were recorded for all single cell line assays, and 30,000 events for experiments involving two cell lines. All data were collected using FACSDiva (BD Biosciences)

### synNotch and SNIPR Activation Assays

Jurkats expressing synNotch and SNIPR constructs were seeded at a density of 2.5e4 cells/well in a U bottom 96 well plate. Cells were plated alone or with an equal number of target K562 cells in a total of 200µl of growth medium. Plates were spun briefly (400xg for 1 minute) to increase likelihood of cell-cell interaction. Cells were co-cultured for between 24 and 72 hours depending on the assay, and the BFP expression and surface expression of receptors and Spike were conducted using flow cytometry. SynNotch and SNIPR activation was assessed as described previously by fitting a two-component Gaussian mixture model to BFP expression data and estimating the fraction of the population in the ‘off’ and ‘on’ components^18^.

### CAR Evaluation Assays

Target K562 cells were spun down, resuspended in T cell growth medium, and were plated in U bottom 96 well dishes at different densities using serial dilution with the 2.5e4 cells plated in wells for the 1:1 effector:target condition. CAR-expressing T cells were stained 1:10000 with CellTrace CFSE (ThermoFisher #C34570) for 20 minutes at 37C and then washed with DPBS. Stained cells were then plated at a density of 2.5e4 cells per well with and without target cells. Cells were co-cultured between 24 and 72 hours depending on experimental conditions. At the selected endpoint, cells were pelleted, stained for either CD25 expression in co-culture conditions or surface markers (anti-Myc or anti-CD19) in mono-culture wells. Cells were then assessed via flow cytometry. T cells and K562s were separated by gating on SSC and CellTrace fluorescence, with K562s having low CellTrace fluorescence and moderate to high SSC values and T cells having low SSC values and moderate to high CellTrace fluorescence. To calculate % target lysis, the number of K562 cells was averaged for each K562 target line across the three replicate wells where those targets were plated with untransduced T cells and then the number of K562s in all experimental wells for that cell line was divided by that number.

### SARS-CoV-2 culture

SARS-CoV-2 isolates USA-WA1/2020 (BEI NR-52281), California (B.1.429), Beta (B.1.351, California Department of Health), Delta (B.1.617.2, California Department of Health), and UK (B.1.1.7, California Department of Health) was used for infection studies. All virus experiments were performed in a Biosafety Level 3 laboratory. SARS-CoV-2 stocks were propagated in Vero-TMPRSS2 cells and their sequence verified by next-generation sequencing. Viral stock titer was calculated using plaque-forming assays.

### Viral Infection Studies

Jurkat cells were seeded into 96-well plates at 2.5e4 cells per well. Cells were rested for at least 24h before infection. At the time of infection, medium containing viral inoculum (MOI of 0.1 and 1) was added on the cells. At 72 hours post infection the cells were washed once with PBS-/- and then incubated for 30 minutes at 4C with 1:1000 Bioscience Fixable Viability Dye eFluor 780 (Invitrogen #65-0865-14) in PBS-/-before fixing with 4% PFA for 30 minutes, washed twice with PBS-/-, and stored at 4C for further processing. Cells were prepared for flow cytometry by pelleting, resuspending in flow buffer, and then were assessed for BFP expression in cells without uptake of the viability dye. Percent population activation was assessed as described above in synNotch and SNIPR assays.

### Plaque Forming assay

Cell supernatants were analyzed for viral particle formation for viral stocks. In brief, Vero-TMPRSS2 cells were seeded and incubated overnight. Cells were inoculated with 10−1 to 10−6 dilutions of the respective homogenates or supernatant in serum-free DMEM. After the 1-h absorption period, the media in the wells were overlaid with 2.5% Avicel (RC-591, Dupont). After 72 h, the overlay was removed, and the cells were fixed in 10% formalin for 1 h and stained with crystal violet for visualization.

### Data Presentation, Analysis, and Availability

All experiments were performed in at least biological triplicate. All data bars are the mean of 3 replicates, with error presented as standard deviation of replicates, and individual replicate values presented as circles where possible. For flow cytometry experiments, fluorescence is reported as median fluorescence value for all cells within a single replicate, and the presented means and error are calculated between replicates. Events collected from flow cytometry were filtered to remove small events and then gated on FSC and SSC to capture singlet populations.

Where presented, histogram density estimates represent all events for all 3 replicates in a flow cytometry experiment, and are calculated via seaborn’s kdeplot function with bandwidth adjustment of 0.2. All data analysis was conducted using custom Python scripts, available on github^76^. Analysis was conducted in Jupyter^77^ and relied on numpy^78^, matplotlib^79,80^, seaborn^81^, pandas^82,83^, SciPy^84^, scikit-learn^85^ and fcsparser. Protein schematics were inspired by their respective structures as indicated in figure legends and assembled used pieces from a proposed protein emoticon^86^. All primary data from flow cytometry experiments are available on Zenodo^87^. All figures assembled with Affinity Designer 2.

## Supporting information

Supplemental Table 1 - Constructs

DNA Sequences

## Acknowledgments

We thank the members of the Hana El-Samad, Wendell Lim, and Jeremy Reiter laboratories for their comments and feedback throughout this project, especially Hersh Bhargava. We thank Manny De Vera, Pilar Lopez, and Sim Sidhu for essential assistance. Human T cells used in this study were generously isolated and donated by the Lim lab. We thank the Whelan laboratory for providing the Vero cells overexpressing human TMPRSS2 (Vero-TMPRSS2). ZYW thanks Bad Moves, Martha, Stephanie E. Crilly, and Kayleigh S. Stevenson for essential support.

## Author Contributions

ZYW: Conceptualization, formal analysis, funding acquisition, investigation, methodology, resources, writing - original draft, writing - review & editing SS: Investigation, methodology, resources, formal analysis, writing - original draft MK: Investigation, methodology, resources, writing - review & editing DHS: Investigation, methodology, resources, writing - review & editing IPC: Investigation, methodology, funding acquisition, writing - original draft, writing - review & editing MO: Supervision, funding acquisition, resources WAL: Supervision, funding acquisition, resources HE-S: Conceptualization, supervision, funding acquisition, writing - review & editing

