## Supplemental Figures for "*De novo*-designed minibinders expand the synthetic biology sensing repertoire"

### Supplemental Figures for Weinberg, Soliman et al. 2024

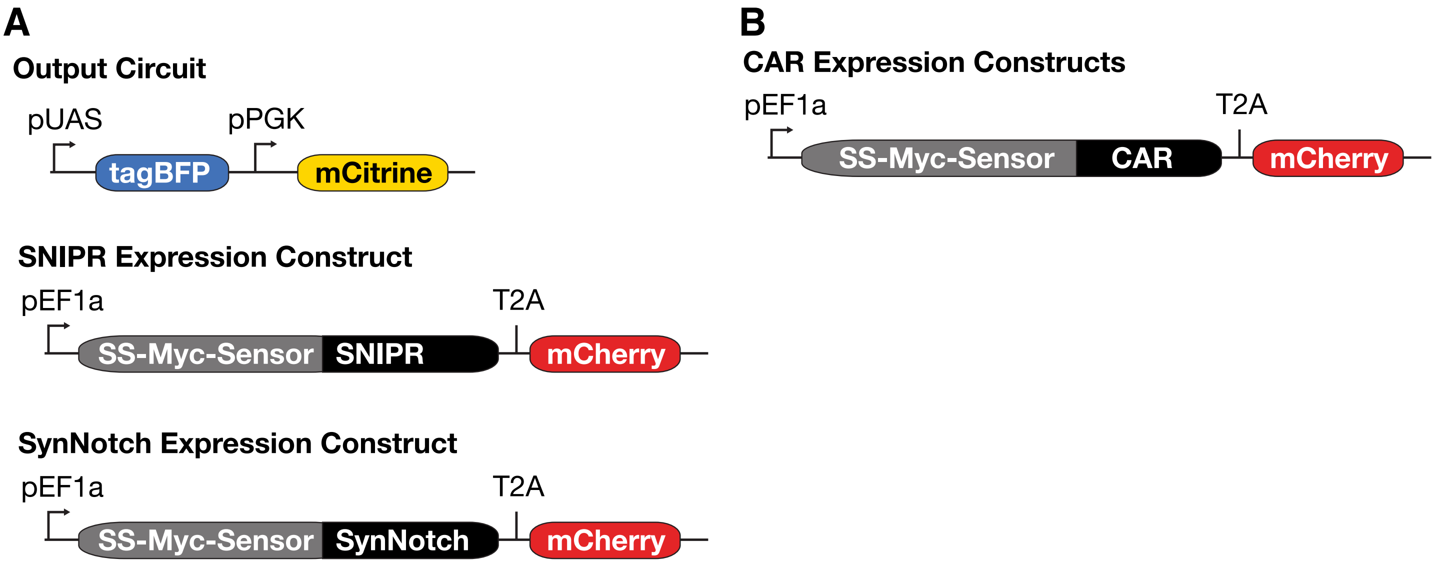


**Supplemental Figure 1:** Expression construct diagrams. A) Constructs used in SNIPR and synNotch experiments in Jurkats. SS = CD8a signal sequence, Myc = Myc epitope tag. B) Construct used in CAR experiments. Figure adapted from Weinberg et al. 2021.

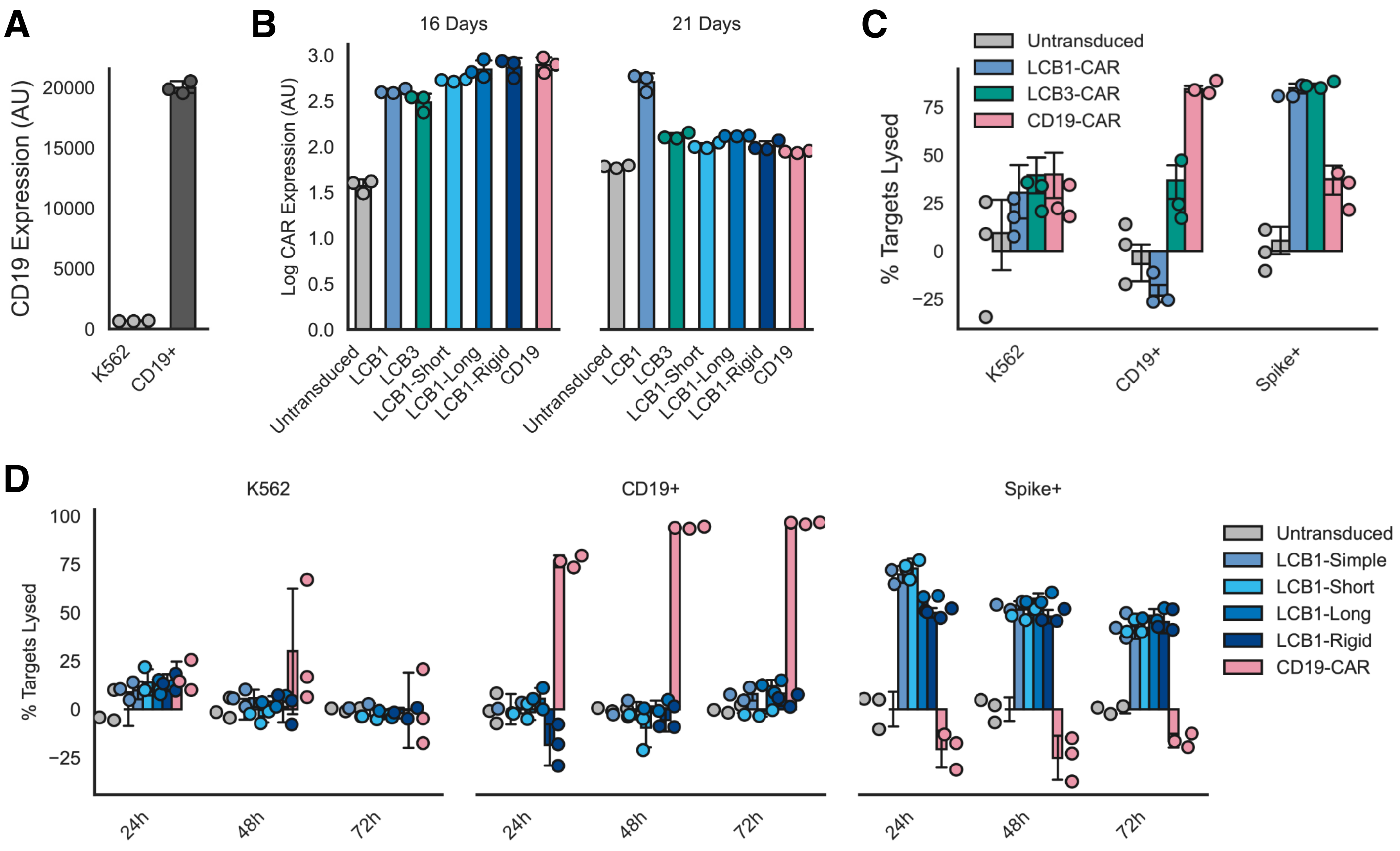


**Supplemental Figure 2: Further characterization of minibinder CAR.** A) Expression of CD19 on K562 targets as assessed by anti-CD19 antibody staining ( (median fluorescent counts for populations across 3 replicates). B) Expression of CAR variants as assessed by anti-Myc staining of extracellular Myc epitope on each receptor after 16 days (initial expression test) and 21 days (completion of killing experiment) (median fluorescent counts for populations across 3 replicates). C) Comparison of CAR function in T cells from a different human donor than Figure 5A&C. Fraction of targets lysed for each of 3 K562 lines across both minibinder receptors and the CD19 benchmark receptor. D) Fraction of targets lysed as a function of time for each LCB1 linker variant compared to CD19-CAR in T cells from the same human donor as Figure 5A&C. All error bars are standard deviation.

**A**

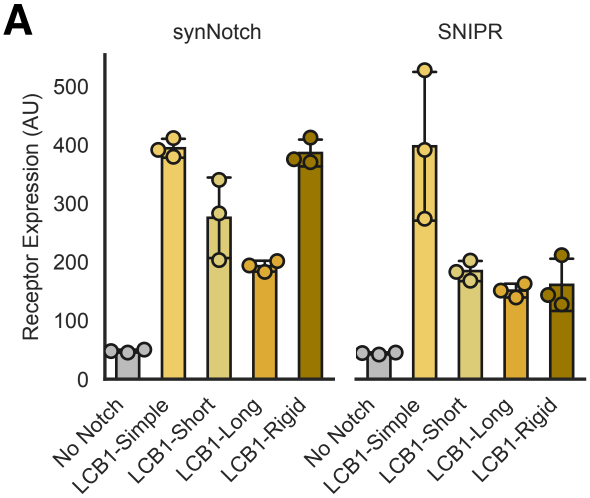


**Supplemental Figure 3: Receptor expression for LCB proteolytic receptors with linker variants.** A) Receptor expression as assessed by anti-Myc staining of an extracellular Myc epitope on each receptor for synNotch and SNIPR receptors with 4 linker variants and the LCB1 minibinder (median fluorescent counts for populations across 3 replicates). Error bars are standard deviation.

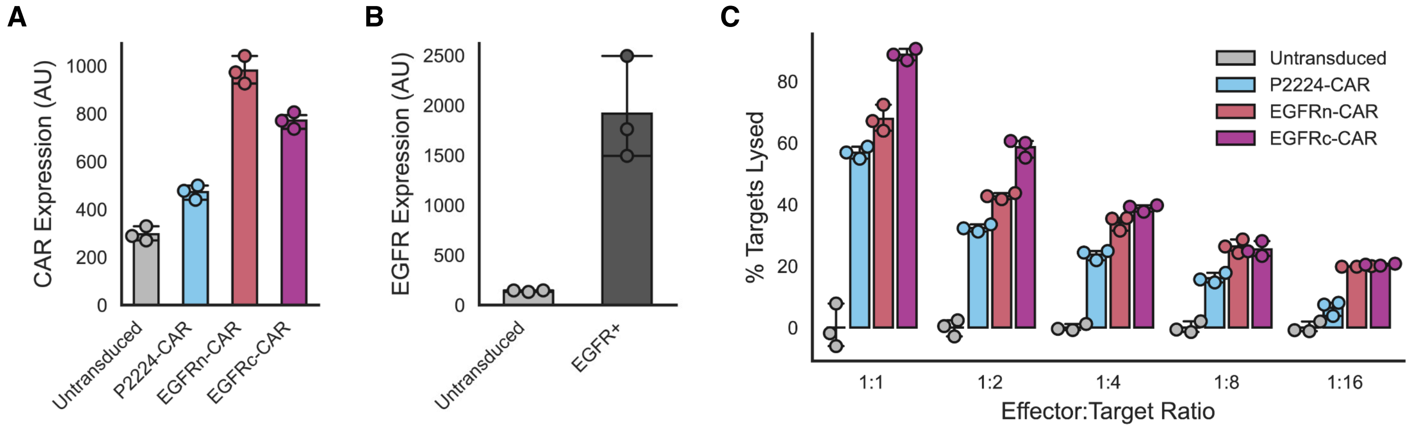


**Supplemental Figure 4: Further characterization of anti-EGFR CARs.** A) Expression of CAR variants as assessed by anti-Myc staining of extracellular Myc epitope on each receptor after 14 days. B) Expression of EGFR antigen on untransduced and EGFR+ K562 cells as assessed by anti-EGFR staining. C) The fraction of target cell populations that were lysed after T cell incubation as a function of the ratio of effector T cells to K562 targets that were incubated together. All error bars are standard deviation.
